# ‘Switch-like’ transition from random to directed motility of microtubules by a yeast dynein

**DOI:** 10.1101/181404

**Authors:** K. Jain, N. Khetan, C. A. Athale

## Abstract

Processive transport by multiple molecular motors that step stochastically, requires a form of mechanical coupling. In a quantitative microtubule (MT) gliding assay with yeast cytoplasmic dynein, we investigate the nature of this coupling by examining the effect of MT length and motor density on transport. We find speed and velocity have a length dependence for low motor numbers, but are independent of MT length for high motor densities. The dependence of speed, velocity and degree of randomness of MT transport is best understood when evaluated in terms of the numbers of motors bound to a filament. A model of collective transport of MTs, based on stochastic stepping and asymmetric detachment rates, reproduces the experimental trends of decreasing diffusivity with increasing number of motors. Additionally, the model predicts a ‘switch-like’ increase in directionality of MT transport above a threshold number of motors. Such a rapid transition from random to directed motility with increasing numbers of yeast dyneins, could play a role *in vivo* during mitosis in the ‘search and orientation’ of the *Saccharomyces cerevisiae* nucleus.

## Introduction

Microtubule (MT) based molecular motors play a vital role in regulating cell physiology and mechanics. The transport of MTs by motors is essential in eukaryotic cell division (1, 2), polarization (3) and migration (4). Single-molecule *in vitro* assays have increasingly provided a detailed understanding of the mechanochemistry of kinesins (5–7) and dyneins (8–10). However, *in vivo* scenarios typically involve multiple motors acting together, as seen in vesicle transport (11), spindle positioning (12) and assembly (13) and the development of neuronal axons (14, 15).

The study of dyneins at a single molecule level has gained momentum more recently owing to their complexity (16), in contrast to kinesins, which have been better studied (5, 17, 18). The stepping behavior of dyneins varies depending on the source, as seen in the load dependence of step sizes in mammalian (8) and *Dicytostelium* (11) dyneins. In contrast, ciliary dyneins from *Tetrahymena* (19) and cytoplasmic dyneins from *Saccharomyces cerevisiae* (9) take discrete 8 nm forward steps. Opposing loads on *S. cerevisiae* dyneins result in probabilistic backward stepping (20, 21). While a small number (up to 7) of these dyneins are reported to be processive (22), *in vivo* multiple copies of this dynein slide MTs and position the nucleus during cell division (23, 24). A better understanding of the quantitative nature of linear MT transport by multiple dynein molecules, could improve our understanding of the role of linear filaments bound by increasing numbers of motors in nuclear positioning during mitosis.

Such multi-motor transport scenarios have also been addressed using theoretical models such as the transport of vesicles (25–27) and linear MT filaments, i.e. gliding assays (28, 29). Some of these models have also been successfully reconciled with experiments as in the case of gliding assays of actin transport by myosins (30) and MT transport by kinesins (29, 31). These models have been used to explain the spontaneous emergence of spirals (30), the co-existence of multiple velocities at the same motor density (32, 33) and the resolution of tug-of-wars in aster transport (34). These ‘emergent’ properties result from multi-motor coupling, and emphasize the need to better understand the theoretical basis for collective transport by reconciling to experiment.

While individually both kinesins and dyneins step stochastically (9, 35, 36), the collective behaviour of both motor types involves a form of cooperativity. This is evidenced by the increase and saturation in velocity with increasing kinesin density (5). A model of ‘loose coupling’ of motors (37, 38) has been observed in kinesin-1 (39), while other mechanisms such as fluctuation based load-sharing (35, 40) and negative cooperativity of detachment (41, 42) have also been proposed to enable collective transport. Recently asymmetric detachment has also been shown in single molecule experiments to be involved in cooperative transport by the *S. cerevisiae* cytoplasmic dynein (43), and predicted in simulations to increase cargo travel distances (44). However, the consequences of these properties on collective transport by large numbers of motors are yet to be experimentally tested.

Here, we examine the *in vitro* transport of linear MTs by the minimal *S. cerevisiae* dynein, that has served as a model dynein in single-molecule studies (9, 20, 21, 43) and compare it to simulations. We reconstitute a gliding assay based on this dynein and use calibrated motor density and the intrinsic variability of MT lengths to examine the quantitative effects of motor numbers on filament transport. In parallel, we have developed a theoretical model of the single motor mechanics, to simulate a 2D gliding assay with MT mechanics and thermal effects, in order to compare theory with experiment (Figure 1). Based on the measures of speed, directional velocity and diffusivity of transport, as a function of motor density and MT lengths, we infer the potential functional relevance of such collective transport in yeast nuclear positioning during mitosis.

**Figure 1.**
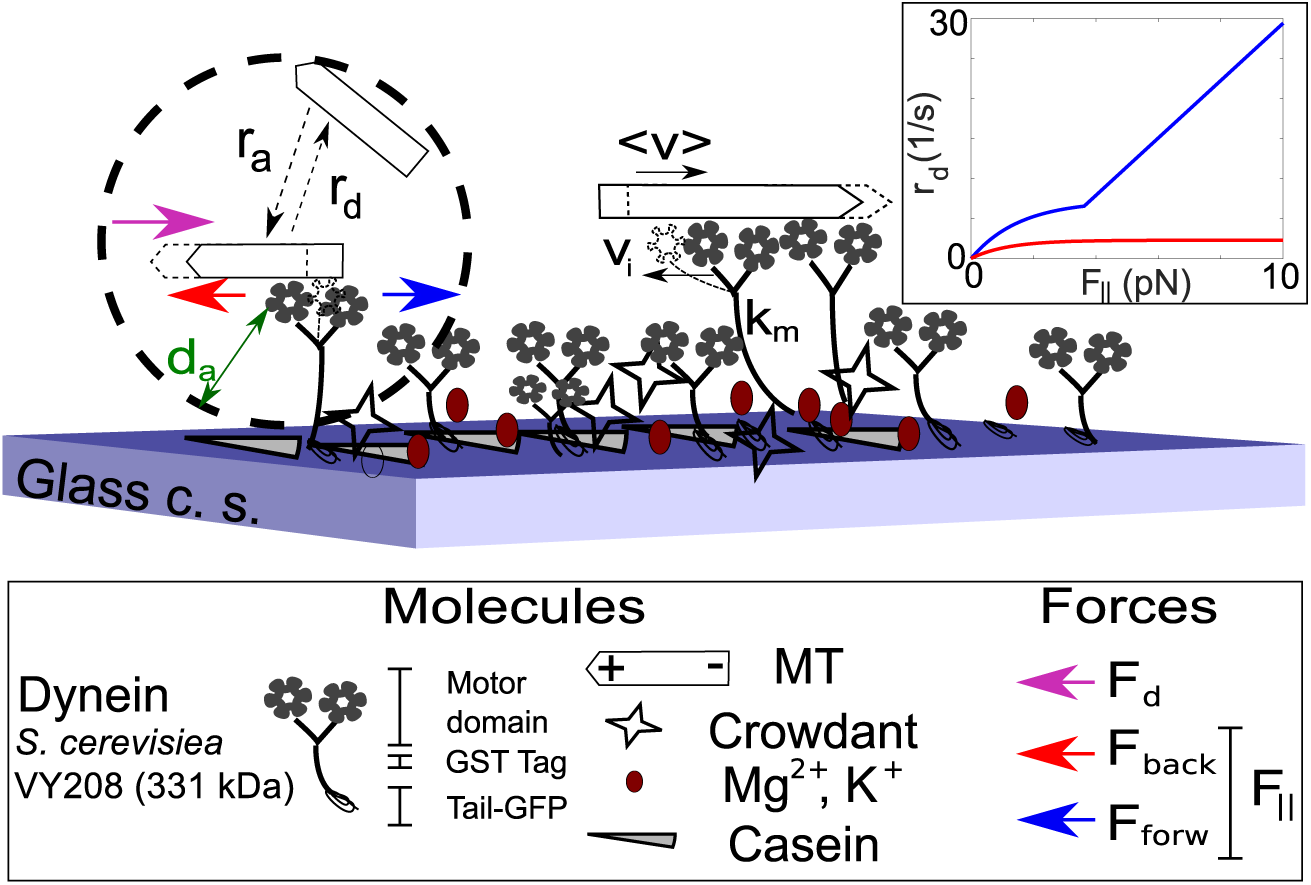
Dynein gliding assay in experiment and simulation. The molecular components of the *in vitro* reconstituted MT-transport by surface immobilized dynein motors in experiment are schematically represented. A stochastic mechano-chemical model of the system is developed in parallel, where motors attach to MTs at a rate *r*_*a*_ if they are within an attachment distance *d*_*a*_, and detach at a rate *r*_*d*_. Single motors step along the MT with an instantaneous velocity *v*_*i*_ and multiple motors transport MTs with an effective velocity *(v)* (black arrows represent the direction of movement). A drag force *F*_*d*_ (magenta arrow) acts on the MTs due to their motion through the medium with viscosity *η*. The motors are modeled as Hookean springs with a stiffness *k*_*m*_. Motor stepping along MTs produces a parallel force *F*_*||*_ in forward (blue arrow) and backward (red arrow) directions, affecting the motor detachment rate (inset).

## Results

### Gliding assay in experiment and simulation

We have used a combination of experiment and theory to examine if increasing numbers of a truncated minimal dynein motor are ‘coupled’ or interfere with one another in collective transport. To this end, we performed a quantitative gliding assay using a minimal, truncated *S. cerevisiae* cytoplasmic non-essential dynein (Dyn 331 kDa) that is comparable in processivity to the full-length motor (9). This dynein with an intact motor domain, a GST-domain for dimerization and a GFP-tagged tail but missing segments of the linker (Figure 1), has become a model for *in vitro* testing of dynein-based transport (21, 22). The motor is immobilized either non-specifically or by binding to immobilized anti-GFP antibodies and after blocking, taxol stabilized MTs are imaged. Simultaneously, we have developed a model to simulate MT transport by immobilized motors for comparison with experiment. The model includes single motor velocity (*v*_*i*_), spring constant of motors (*k*_*m*_), attachment (*r*_*a*_) and detachment rates (*r*_*d*_) that determine whether the MT is bound to motors and the role of forces acting on the motor in the forward (*F*_*forw*_) and backward (*F*_*back*_) directions. Along with motor generated forces, thermal forces and the drag force acting on MTs (*F*_*drag*_), all collectively result in the transport of individual filaments with a mean effective velocity, < *v* > (Figure 1). The model is described in greater detail in the Materials and Methods section. We began by examining the effect of MT length and motor density on collective dynein transport.

### Degree of randomness of MT transport in experiment affected by motor density and MT lengths

The experimental time-series of gliding MTs were tracked to quantify single filament motility with a motor density of 2 motors/*μ*m^2^ (Figure 2(A), Video 1 (A)) and 16 motors/*μ*m^2^ (Figure 2(B), Video 1 (B)) using a nanometer precision filament tracking program, FIESTA, which uses an extended Gaussian PSF model for super-resolution tracking (45). Motor densities were estimated from a calibration of the PSF (Figure S1 (A)-(C)) and mean intensities of increasing concentrations of GFP (Figure 1(D)) to determine the concentration of GFP-dynein on the surface. Motor activity was confirmed using a phosphate release assay, compared to controls (Figure S2). The lengths of the MTs are exponentially distributed with mean lengths 4.81 *μ*m and 2.33 *μ*m, before and after shearing respectively (Figure S3). This length distribution combined with two different motor densities, allowed us to examine a wide-range of motor numbers and their effect on MT transport.

**Figure 2.**
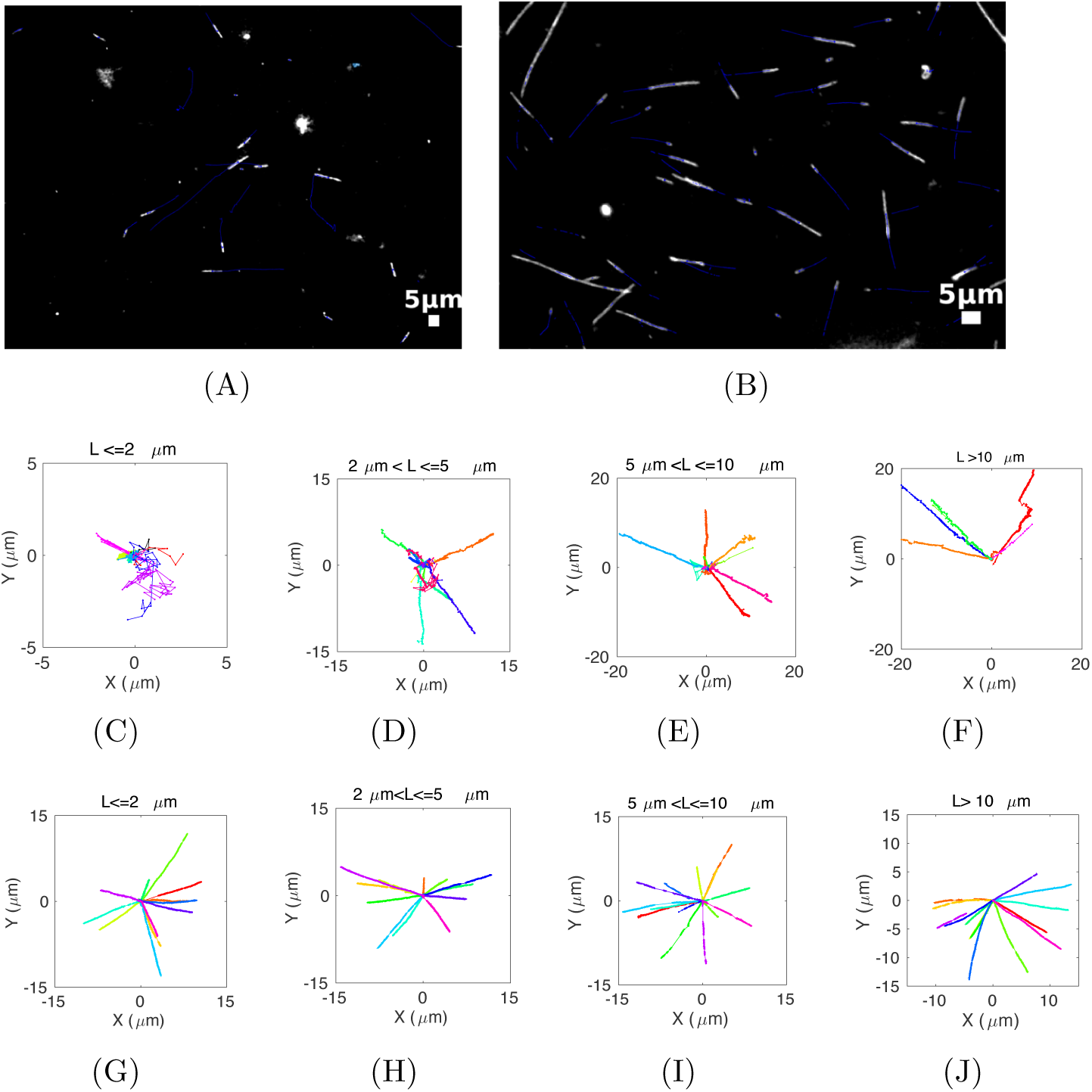
Microscopy of gliding MTs. (A), (B) The first frame of a time-series of gliding MTs (Video 1(A), (B)) labelled with rhodamine-tubulin (white) in the presence of surface immobilized yeast dynein of density (A) *ρ*_*m*_= 2 and (B) *ρ*_*m*_= 16 motors/*μ*m^2^ is overlaid with a time-trace of MT centroids (blue) obtained from single filament tracking. (C)-(J) The corresponding representative xy-trajectories of the leading ends of MT filaments of length ranges (C),(G) L*<* 2, (D),(H) 2 *<*L *≤* 5, (E),(I) 5 *<*L *≤* 10, (F), (J) L*>* 10 *μ*m are compared between experiments with motor densities (C)-(F) *ρ*_*m*_= 2 and (G)-(J) *ρ*_*m*_= 16 motors/*μ*m^2^. The colors represent tracks of individual filaments (n= 5 to 15).

At a low motor density (2 motors/*μ*m^2^) the XY trajectories of MT filaments appear random for filaments of lengths less than 2 *μ*m (Figure 2(C)) and become increasingly directional for longer MTs ranging in lengths between 2 to 5 *μ*m (Figure 2(D)), 5 to 10 *μ*m (Figure 2(E)) and greater than 10 *μ*m (Figure 2(F)). In contrast, the qualitative trends of XY trajectories at a higher motor density (16 motors/*μ*m^2^) appear unchanged, independent of MT lengths (Figure 2(G)-2(J)). The leading end of MT filaments were plotted in all cases for consistency, but qualitatively similar effects were observed, when the trailing ends were plotted. Filament centroids were not used due to the loss of information about potential swiveling motion. This suggests both length and motor density determine whether MT transport will be random or directed. We proceeded to examine if a similar qualitative trend was to be seen in simulations with a detailed single-motor mechanics model based on experimentally measured properties of dynein.

### Simulated MTs qualitatively transition from random to directed transport with increasing length and motor density

We have modeled MT gliding on a sheet of uniformly distributed immobilized dynein motors that are modeled as stochastic discrete steppers with attachment and detachment dynamics, as described in detail in the Model section. The simulations outputs, which depict only bound motors for clarity, suggest MT transport is more random with frequent changes of direction at a motor density *ρ*_*m*_ = 2 motors/*μ*m^2^ (Figure 3(A), Video 2(A)) as compared to *ρ*_*m*_ = 16 motors/*μ*m^2^ (Figure 3(B), Video 2(B)). Increasing lengths of MTs transported by *ρ*_*m*_ = 2 motors/*μ*m^2^ transition from an apparent random walk for short filaments*L* = 1 *μ*m (Figure 3 (C)) and *L* = 4 *μ*m (Figure 3 (D)), to relatively more directed motility for long filaments*L* = 7 *μ*m (Figure 3 (E)) and *L* = 10 *μ*m (Figure 3 (F)). At a higher motor density, this transition is more marked for the same length range of *L* = 1 to 10 *μ*m (Figure 3 (G)-(J)). This result suggests both MT length and motor density qualitatively affect motility, similar to our experimental observations. However in simulations, we observe a length-dependence of the qualitative behavior of filament transport at both low and high motor densities, an effect not seen in experiment. In order to more precisely compare experiments and simulations, we proceed to quantify motility statistics.

**Figure 3.**
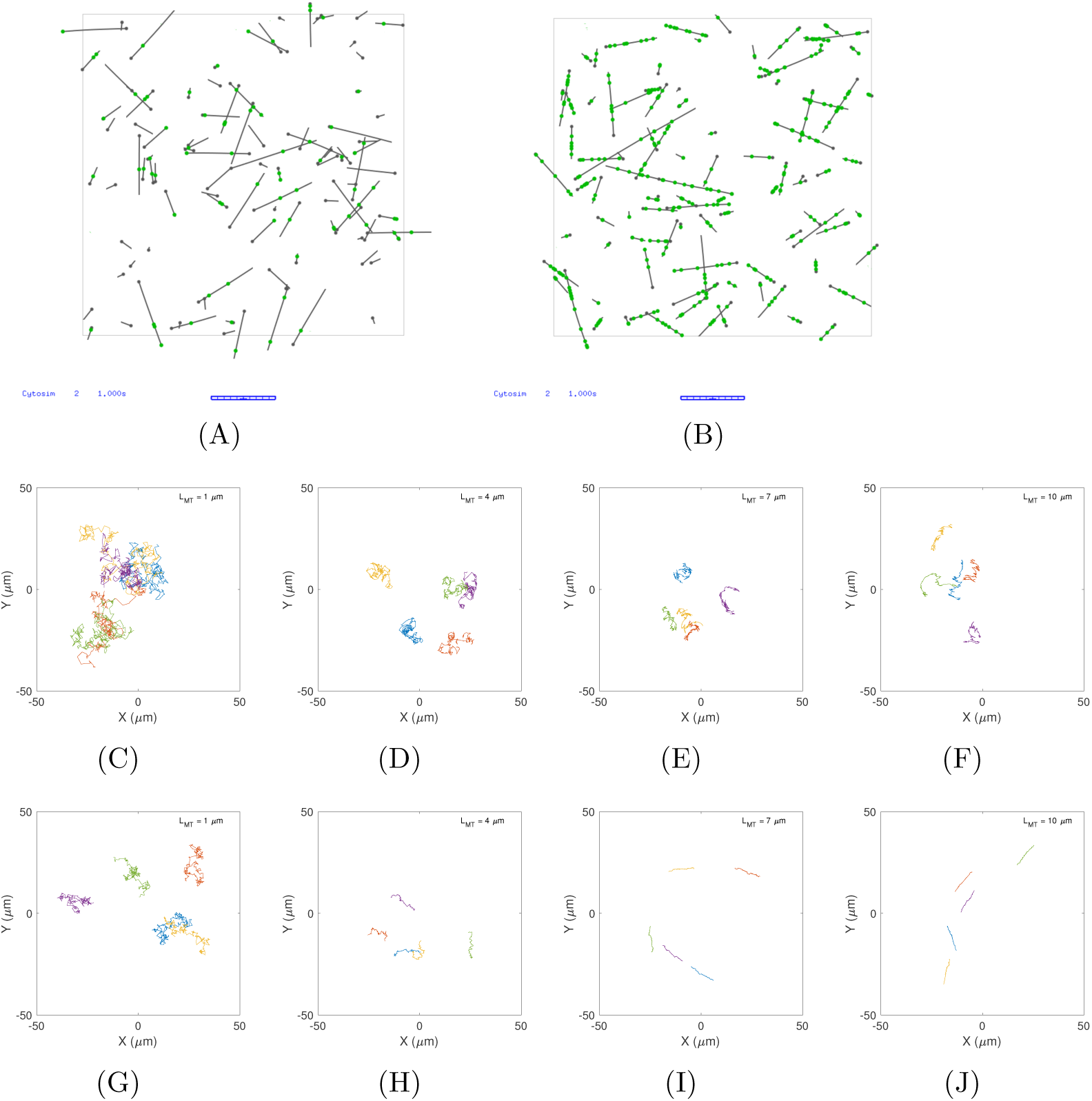
Simulation of gliding MTs. (A), (B) The first frame from simulations (Video 2(A),(B)) of gliding MT filaments (grey lines) in the presence of immobilized dynein motors (green filled circles) is visualized for motor densities of (A) *ρ*_*m*_= 2 and (B) *ρ*_*m*_= 16 motors/*μ*m^2^. MT plus ends are marked by filled grey circles. Only bound motors are displayed for clarity. Scale bar: 10 *μ*m. (C)-(J) The XY trajectories of representative filaments (n=5) of lengths (L) corresponding to 1, 4, 7 and 10 *μ*m are plotted in the presence of density of motors of (C)-(F) 2 and (G)-(J) 16 motors/*μ*m^2^. Colors represent individual tracks.

### Density dependence of speed and velocity of MT transport in experiment and simulation

The experimental data from single-filament tracking was used to estimate MT transport speed statistics (scalar, considering only the magnitude), pooled from multiple fields of view with motor densities of *ρ*_*m*_= 2 motors/*μ*m^2^ (*N*_*MT*_= 229) and *ρ*_*m*_= 16 motors/*μ*m^2^ (*N*_*MT*_ = 1770). Short MTs appear to move faster than long ones at a low motor density, while at a higher motor density speed is unaffected by length (Figure 4(A)). The simulated transport of MTs by the same motor densities shows reasonable agreement with experiment. The higher MT speeds seen in simulations for short filaments at both low and high motor density, as compared to experiment are likely to be due to the extent to which diffusive filaments are not captured in experiment when they leave the imaging plane.

**Figure 4.**
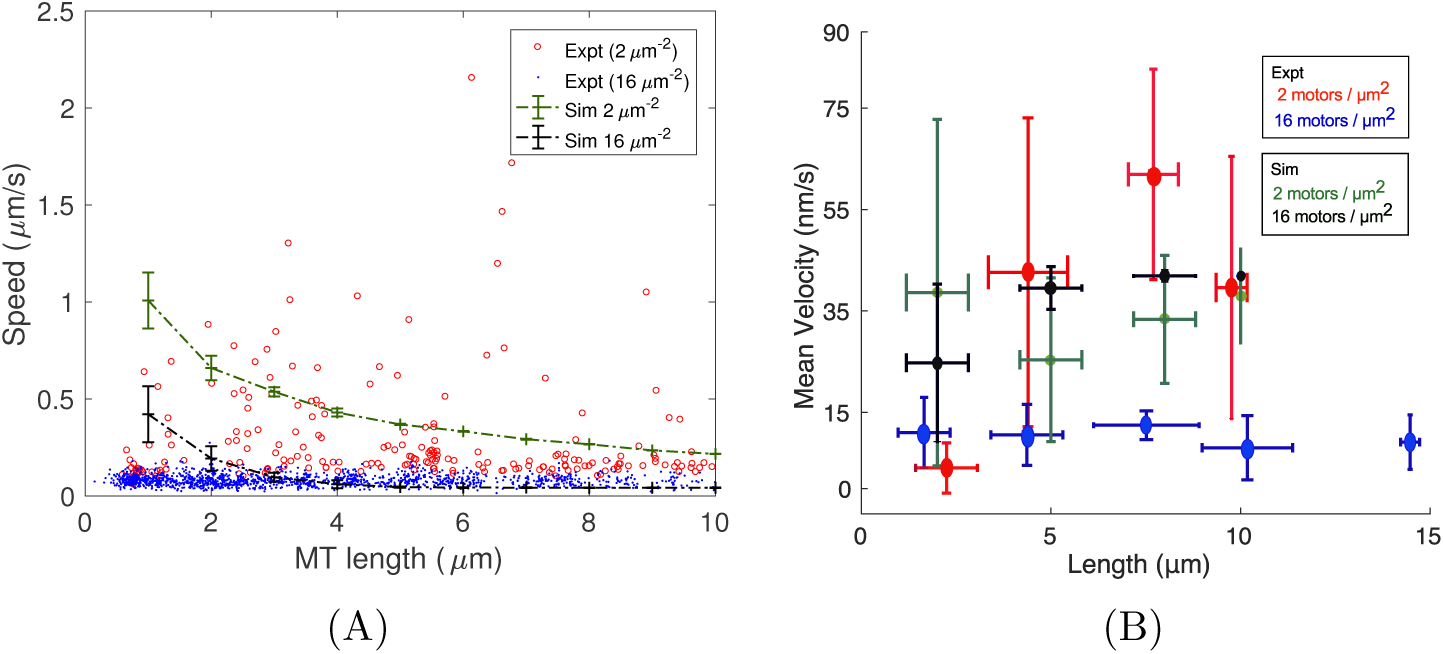
Speed and velocity compared between experiment and simulations. (A) Experimentally measured mean speeds (*μ*m/s) of filament transport with a motor density of *ρ*_*m*_ = 2 (*N*_*MT*_ = 229) (red circles) and *ρ*_*m*_ = 16 motors/*μ*m^2^ (*N*_*MT*_ = 1770) (blue circles) are plotted as a function of MT length (*μ*m). MT speeds from simulations (mean*±*s.d., *N*_*MT*_ = 100 for each length) are overlaid for the same densities. green dashed line: *ρ*_*m*_ = 2, black dashed line: *ρ*_*m*_ = 16 motors/*μ*m^2^. (B) The mean velocity as a function of MT length class is plotted from experiments (red, green) and compared to simulations (green, black) for dynein densities *ρ*_*m*_ = 2 (red) and *ρ*_*m*_ = 16 motors/*μ*m^2^ (blue). Only bound filaments were considered for the analysis of simulation data.

The mean velocity (the vector change in position with time) estimated from experiment with 2 motors/*μ*m^2^ increases slightly with MT length, while in presence of 16 motors/*μ*m^2^ appears unchanged (Figure 4(B)). In simulation, velocities appear to slightly increase with MT length, for both low and high motor densities, suggesting a slight increase in directionality with length.

This suggests the high speed of transport of short filaments at a low motor density is the net result of motor activity and diffusion, potentially due to a lower probability of a filament being bound. On the other hand, long filaments and high motor densities result in processive transport. To better understand the degree of diffusivity of short filaments and the processive transport of long filaments by apparent collective mechanical coordination, we proceeded to estimate the effect of absolute numbers of motors on gliding MTs from experiment and simulation.

### ‘Switch-like’ transition in drift velocity and directionality of MT transport with motors

The proportion of effective diffusion and drift velocity contributing to MT transport was estimated from MSD plots of experimental trajectories by fitting them to equations that represent (a) anomalous diffusion (Equation 2) and (b) diffusion with drift (Equation 3). All the plots of MSD for increasing MT lengths (representative plots of MT lengths, *L*_*MT*_ 1.48 to 15.5 *μ*m) from experiment (Figure S4 (A)) indicate super-diffusive transport, based on the value of the anomaly parameter *α ≥* 1.5 obtained by fitting the anomalous diffusion model (Figure S5 (A), (B)). The MSD from simulations of increasing MT lengths *L*_*MT*_ = 2, 5, 8 and 10 *μ*m (Figure S4 (B)) results in *α* increasing from 1 to 2, i.e. transitioning from diffusive to super-diffusive transport (Figure S5 (C), (D)). The fit to the diffusion with drift model (Equation 3) demonstrates the experimental estimate of the effective diffusion coefficient (*D*_*eff*_) is high in presence of a few motors (less than 10), and low for motor numbers exceeding 10 (Figure 5(A)). The median drift velocity (*v*_*drift*_) values increase with number of motors up to a threshold (15 motors), but then decrease for motor numbers that exceed this threshold (Figure 5(B)). In simulations *D*_*eff*_ is either constant or decreases with MT length, depending on the motor density (Figure 6(A)). The same data plotted as a function of motors, collapses onto a single trend of decreasing *D*_*eff*_ (Figure 6(B)). As a function of length, the change of *v*_*drift*_ also varies, depending on motor density (Figure 6(C)). Plotted as a function of motors however, it steeply transitions at ~10 motors from a low (∼ 0 to ∼10 nm/s) to a high value (∼ 40 nm/s) (Figure 6(D)). Consistent with this transition, the anomaly parameter of diffusion (*α*) estimated from simulations, also shows a similar sharp-transition above 10 motors (Figure 5(C)).

**Figure 5.**
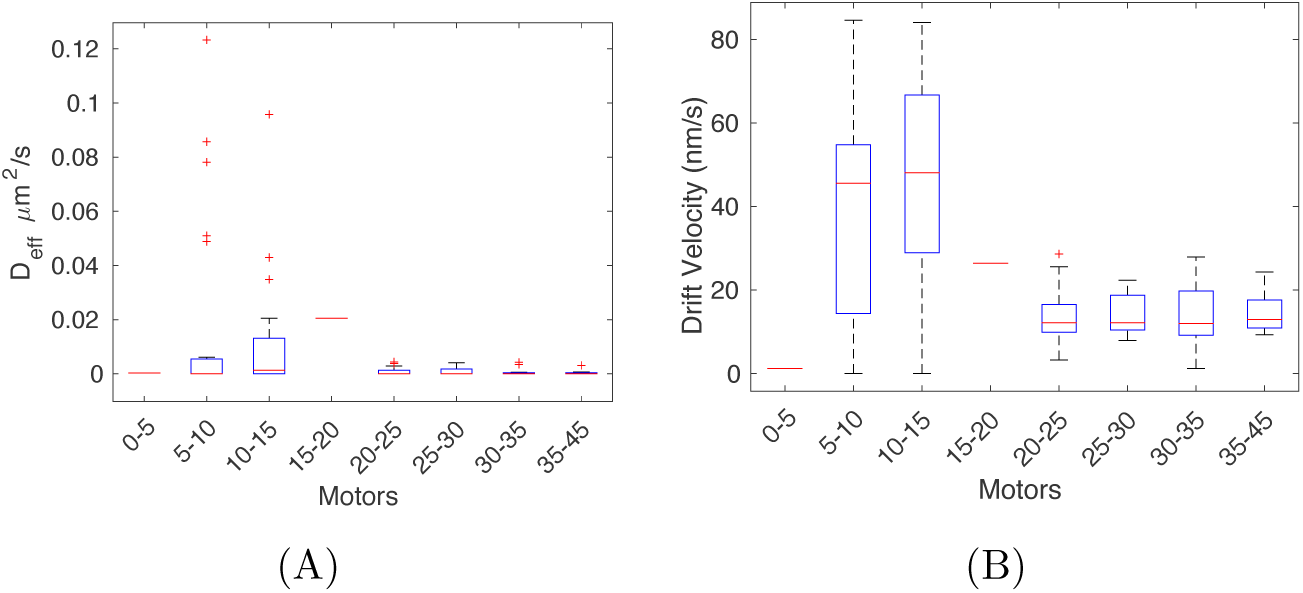
Experimental estimates of diffusion and drift. Fits of the diffusion with drift model to MSD plots were used to infer (A) the effective diffusion coefficient (*D*_*eff*_) and (B) drift velocity (*v*_*drift*_), and are plotted as a function of motor numbers in box plots (red line: median, boxes: second and third quartile boundaries, plus signs: outliers).

**Figure 6.**
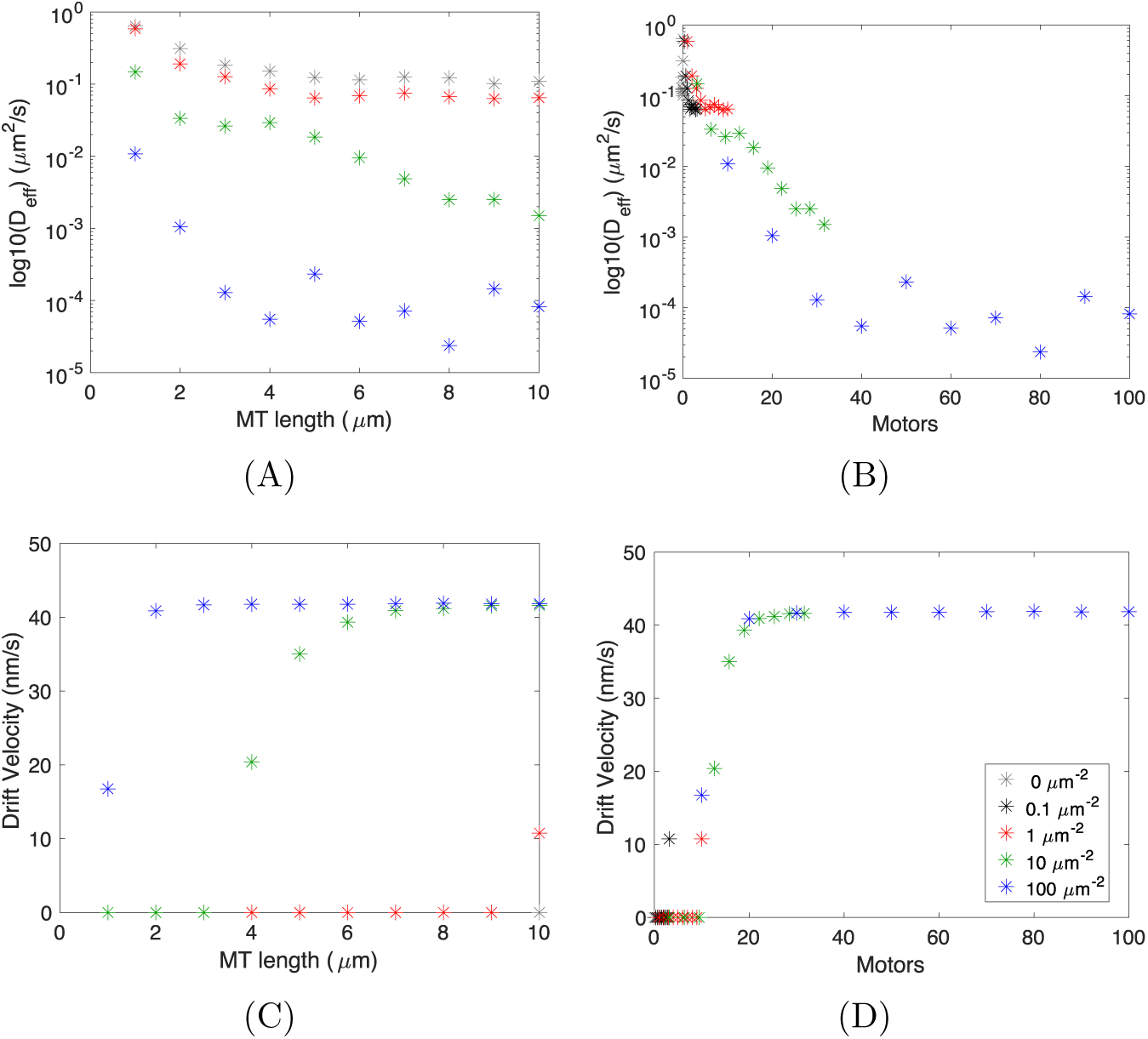
Estimates of effective diffusion and drift from simulation. (A), (B) The effective diffusion coefficient obtained from fitting the diffusion with drift model to the MSD profile of simulated trajectories at different motor-densities and plotted as a function of (A) MT length and (B) motor numbers. (C), (D) The drift velocity similarly estimated is plotted as a function of (C) MT length and (D) motor number. Colors indicate the motor density per *μ*m^2^.

This suggests that the sharp transition in *v*_*drift*_ from diffusive to directed transport with increasing motors predicted in simulations, is only partially observed in experiment. Since MSD analysis is sensitive to the time of observation and numbers of data points, we also quantify the directionality of transport by tortuosity (*χ*), a variable that quantifies randomness in diffusion and drift processes (46). We find experimental values of *χ* increase above ∼10 motors (Figure 7(A)), while simulations demonstrate a ‘switch-like’ transition at ∼ 10 motors from random to directional transport (Figure 7(B)). Thus, experiment and theory suggest a qualitative switch from random to directed transport by yeast cytoplasmic non-essential dynein, as a function of number of motors acting on a filament.

**Figure 7.**
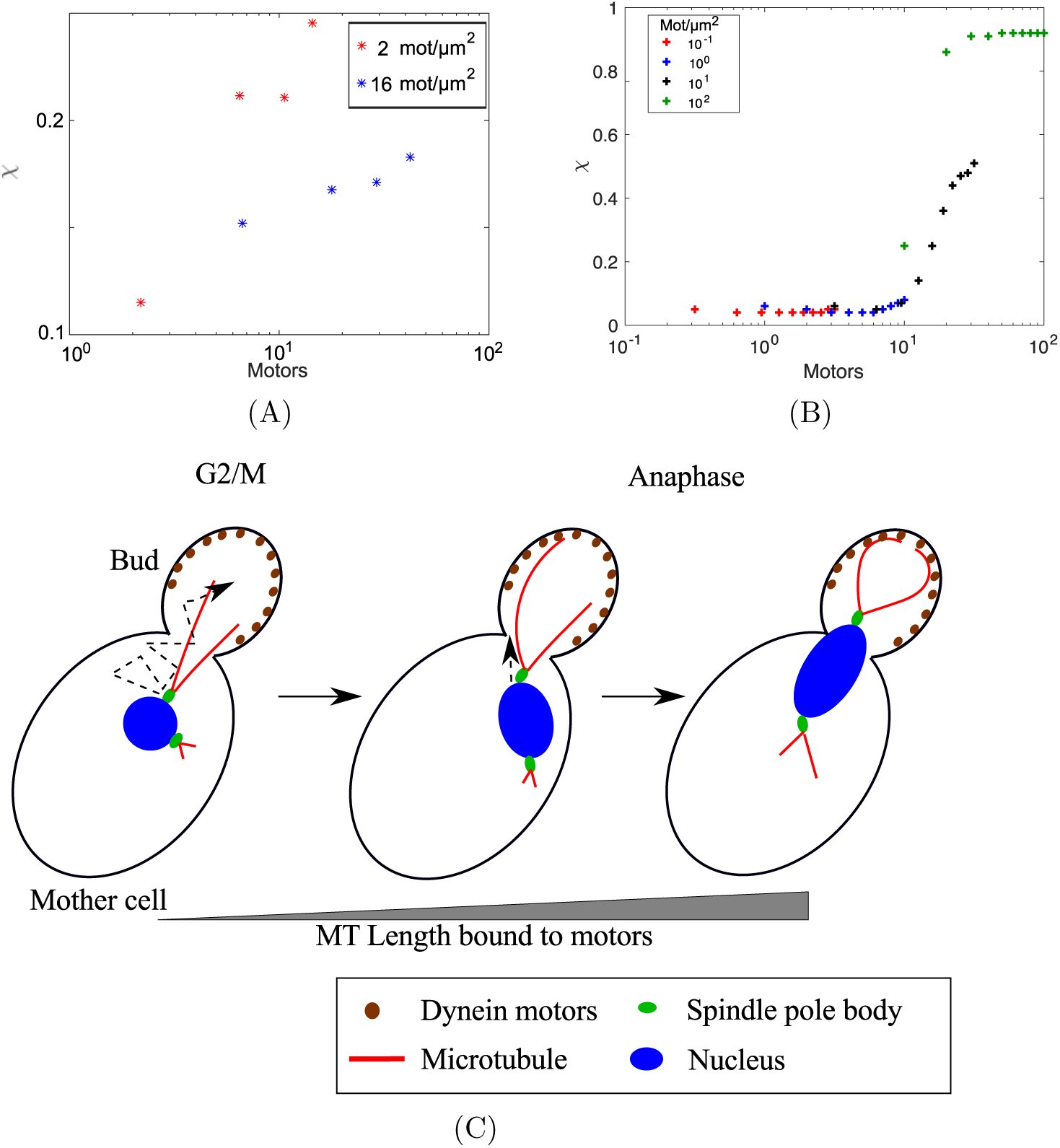
Directionality of MT transport in experiment and simulation. (A) The experimentally estimated median tortuosity (*χ*) values (*N*_*MT*_ *>* 15 for each length) were plotted as a function of motors per filament, using data from motor densities of 2 (red) and 16 motors/*μ*m^2^ (blue). (B) Mean values of *χ* (*N*_*MT*_ = 100 for each length) from simulations, were plotted as a function of motors, combining motor densities of 10^*-*1^ (red), 10^0^ (blue), 10^1^ (black) and 10^2^ (green) and MT lengths ranging from 1 to 10 *μ*m. (C) MT sliding during *S. cerevisiae* mitosis is schematically represented. Astral MTs (red) at the G2/M transition ‘search’ for and are ‘captured’ by cortical dyneins (brown) in the bud cell, followed by ‘reeling in’ and of the nucleus (dashed arrow) resulting in ‘orientation’ in Anaphase along the mother-bud axis – ‘search and orientation’.

## Discussion

In this work, we report collective effects of dynein-based MT transport by comparing quantitative gliding assay experiments with simulations. By varying motor density and using the natural variability of MT lengths in experiment, we find the transport of short filaments is random and of longer filaments is directed, in the presence of a low density of motors. This trend is qualitatively reproduced in simulations. A quantitative comparison of speed and velocity statistics for MTs suggests very short filaments (L < 2 *μ*m) move with a higher speed (scalar) and lower velocity (vector) as compared to long filaments, at a low density of motors. In presence of a high density of motors, both velocity and speed appear to reach constant values. The observed decreasing speed with increasing length, is the result of a decrease in the effective diffusion coefficients *D*_*eff*_, with increasing motor numbers per filament, as expected. The drift velocity, *v*_*drift*_ conversely increases with motors up to a threshold. Simulations predict a ‘switch-like’ transition from random to directed motion above a threshold of 10 motors both in terms of *v*_*drift*_ and tortuosity, *χ*, over a range of 1 to 10^2^ motors per filament. The sudden transition in directionality of transport emerges, we believe, from the single-molecule mechanics of the motor and the mechanical coupling of motors bound to the same filament.

Previous single-molecule studies of yeast dynein have demonstrated the load-dependent, discrete and probabilistic nature of step-size and direction (9, 20). These studies have also identified the ‘minimal’ yeast dynein construct, which we have used in this study to examine it’s collective transport properties. Similar to our results here, the transport of MTs by 1 to 5 molecules of kinesin-1 had revealed a transition from rotational to translational motion with increasing motor number (31). While increasing yeast cytoplasmic dyneins from 1 to 7 had been seen to increase run-lengths of DNA-origami based cargos, a very small difference in the directionality of transport had been reported (22)In our studiy, the MT lengths examined range up to 10 *μm* and longer, resulting in larger numbers of motors (*∼* 100 motors per filament) that could potentially interact with filaments. This could explain the motor-number independence of the directionality of transport seen previously with the same dynein.

The directional persistence of MTs in experiments has been previously evaluated using effective diffusion and drift measures (47, 48). In previous work, we had demonstrated the role of low motor densities on the degree of anomalous diffusion in the transport of MT asters (34). For short filaments at low densities, our results confirm the expectation that speeds of transport of small filaments are more likely to exhibit random motion compared to long filaments, due to the absolute number of motors they encounter (Figure 4(A)). However, at a high motor density (*ρ*_*m*_ > 10^1^ motors/*μ*m^2^), speeds and velocities both remain constant, irrespective of MT length. Consistent with this expectation, the effective diffusion coefficient in experiments decreases with increasing motor numbers. The drift velocity in experiment increases as expected with number of motors and appears to reduce and saturate above a threshold. Simulations reproduce the experimentally observed decrease in *D*_*eff*_ and increase in *v*_*drift*_ with motors. The discrepancy between experiment and simulations in *v*_*drift*_ for the higher number of motors (>10) could be attributed to having fewer samples as compared to simulations. On the other hand, the experimental trend of *χ* appears to qualitatively reproduce the trend seen in simulations (Figure 7 (A),(B)). In future, the predictions of simulations could be tested in experiment, potentially using methods developed recently for micro patterning motors (49, 50). Additionally, the relevance of such a transition from random to directed transport as a function of motor numbers, could also be tested by quantification of the number of motors involved *in vivo* in MT transport.

*In vivo* this cytoplasmic non-essential dynein from *S. cerevisiae* is involved in astral MT sliding during mitosis (23, 51). The role of cortically anchored dynein in both positioning and orienting the nucleus, has revealed a complex network of transport, anchoring and regulatory co-factors (reviewed in (24)). The physical mechanism by which MTs ‘search’ for cortical anchorage points is thought to involve MT pivoting (52) and once ‘captured’, both dynein based pulling and MT shrinkage combine to ‘reel in’ the nucleus and orient it in the bud (23). This scenario observed during G2/M to anaphase transition in *S. cerevisiae* mitosis is reminiscent of a gliding assay with different lengths of MTs binding to immobilized motors and resulting in the sliding of filaments (Figure 7(C)). Our experimental and theoretical observations suggest that the mechanics of the yeast dynein, might be adapted to ensure robust positioning of the nucleus in anaphase. The initial ‘search’ and attachment of filaments to a few motors could provide the ‘noise’ necessary for the nucleus to be correctly oriented along the bud-mother axis. After this ‘search and orientation’ phase, the longer contacts between MTs and motors could then result in the positioning by directed transport, a form of ‘reeling in’. Both careful quantification of the absolute number of motors and the positioning kinetics of the nucleus, could further test our model predictions and the *in vivo* significance of such a mechanism.

In conclusion, we have demonstrated in experiment and simulation with a yeast dynein, the speed of transport decreases and saturates with the number of motors bound to a filament, while the velocity increases and saturates. This trend can be explained by the diffusivity of short filaments due to fluctuations in the number of motors bound. Additionally, the directionality of transport transitions from random to directional with increasing motor numbers above ∼10 motors, an experimentally testable prediction. The model predictions for the *in vitro* scenario, also has implications for the *in vivo* function of dynein in terms of ‘search and orientation’ of the nucleus during mitosis. Thus, we believe the model points to a form of ‘detachment cooperativity’ as a single motor property of the yeast dynein, that could have a functional role in organelle positioning and transport.

## Methods and Materials

### 0.1 Microtubule filament assembly

Rhodamine-labelled bovine (0.8 *μ*g/*μ*l) and unlabelled porcine tubulin (3.3 *μ*g/*μ*l) (Cytoskeleton Inc., Denver, USA) were incubated in BRB80 buffer (80 mM PIPES, 1 mM MgCl_2_, 1 mM EGTA) at 37°C for 15 minutes. BRB80 containing 20 *μ*M taxol was added to obtain fluorescently labelled microtubule filaments. Long filaments (ranging up to 100 *μ*m) were obtained by incubation for ∼ 60 hours. In order to distinguish individual filaments, samples were diluted 10 to 20 fold before microscopy. Short filaments were obtained by shearing the sample using a 20 gauge needle in 4-5 passes.

### Purification of dynein

The truncated minimal dynein construct (331 kDa) from the *S. cerevisiae* VY208 was purified as previously described by (9) and is briefly summarized here. Cells were grown in 2 liters of yeast peptone dextrose (YPD) medium to an optical density at 600 nm of ∼3 and induced with 2% galactose. The culture was centrifuged, lysed and incubated with IgG beads to bind the ZZ-tag of the motor protein. The protein was released from the beads by ∼12 hours treatment with TEV-protease at 4°C, flash frozen and stored at a concentration of 0.06 *μ*g/ml at −80°C. ATPase activity of dynein was assayed in the presence of 20 *μ*l of pre-assembled MT filaments (15 *μ*M goat-brain tubulin, 1 mM GTP, 10% glycerol, 1x BRB80 and incubated for 30 min at 37°C) with 1 mM Mg-ATP (Jena Bioscience, Germany) and malachite green (Sigma, India) in a 96-well plate by measuring absorbance at 660 nm in a plate reader (Varioskan, Thermo Scientific, USA) after 45 minutes and found to be active relative to controls (Figure S2).

### Gliding assay

Flow chambers were prepared by a method similar to that described by (53) using double-backed tape (Venus Traders, Pune) adhered to cleaned microscope slides and 22 x 22 mm coverslips (Micro Aid, Pune, India) to create a chamber with a volume of *≈* 16 *μ*l. Slides and coverslips were cleaned successively in acetone (1 hour), ethanol (15 min), demonized water (twice with for 5 minutes each), 0.1 M KOH (15 minutes) followed by a final rinse with demonized water. The motors were perfused (50 *μ*l of 0.06 *μ*g/ml dynein) through one end of the chamber in lots *sim*15 *μ*l each, incubated for 5 minutes at room temperature. Blocking solution containing 0.5 mg/ml BSA (Sigma Aldrich, Mumbai, India) was flowed in, followed by the addition of 10 *μ*l of MT filaments (labelled, taxol-stabilized) and the slide incubated for 5 minutes. A wash buffer (also used as lysis buffer-30 mM Hepes (pH 7.2), 50 mM potassium acetate, 2 mM magnesium acetate, 1 mM EGTA, 10% glycerol, 1 mM DTT, 1 mg/ml casein) was used to remove unbound motors and MTs. Motility buffer (wash buffer supplemented with 1 mM ATP) with 1x Anti-fade (Cytoskeleton Inc., USA) was perfused to observe MT gliding. For the low density experiments, before adding motors, anti-GFP antibody (0.2 *μ*g /*μ*l) (Sigma Aldrich, Mumbai, India) was perfused through one end of the flow chamber and incubated for 5 mins followed by the addition of blocking solution, motors, MTs, wash- and motility-buffer, as described above.

### Microscopy

The samples were imaged at room temperature (≈ 22°C) using a fluorescent upright microscope, Zeiss Axio Imager Z1 (Carl Zeiss, Germany) with an EC Plan-Neofluar 40x (NA - 0.75) lens, a mercury short arc lamp (X-Cite Series 120, Lumen Dynamics Inc., Canada) and filter sets for rhodamine (520 nm/540 nm excitation and 580 nm/600 nm emission). Images were acquired every second for 10-20 minutes using the Zeiss AxioCam MRm digital camera (Carl Zeiss, Germany) with exposure times 0.7 to 0.9 s.

### Image and data analysis

Image time series of gliding MT filaments were contrast adjusted, denoised using a median filter (radius 2 pixels) and histogram-normalized (0.4% saturated) using ImageJ ver.1.45s (54). The filaments were tracked using a filament tracking tool FIESTA ver.1.04 reported previously (45) in MATLAB R2014b (Mathworks Inc., USA) on a Linux platform. This tool has been widely validated by multiple studies to provide nanometer precision in filament tracking (55–60). The maximal connecting velocity was set to 1 *μm/s* and FWHM to 500 nm, with only those filaments considered for further analysis, that persisted for 10 or more frames and a permitted angular deviation between successive frames of 1 to 5 degrees. Based on the frequency distribution of the change in MT filament lengths over time (dL/dt), a Gaussian function fit with zero mean was used to estimate *σ* and used as a cutoff of change of lengths in further analysis (Figure S7). To ensure that our imaging system was stable, we also tracked a time-series of fixed fluorescently labelled beads (Figure S8(A)). The histogram of the pairwise displacement of beads was used to estimate the modal value of 4 nm as the displacements due to the stage stability (Figure S8(B)), used as a lower limit for MT analysis. From the tracking analysis of MTs, the frequency distribution of MT lengths (L) was obtained and fit to the equation:

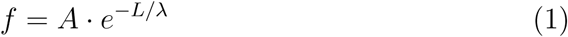

where A is the scaling factor, L is MT-length and λ is the mean length of the population. This was used to distinguish between long (Figure 3(A)) and short filaments (Figure 3(B)). All further data analysis was performed using MATLAB R2014b (Mathworks Inc., USA). The mean speed of MT filaments was estimated from the change in the position of the filament tips at each successive time point, while the mean velocity was estimated from the change in position vectors of successive time points.

### MSD and diffusivity analysis

Using a standard method of MSD calculations as previously described (46), we fit two models of effective diffusion to the 2D experimental and simulated MSD plots: *(a)* anomalous diffusion and *(b)* diffusion with drift. The *(a)* anomalous diffusion model is:

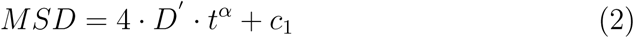

where *D’* is the anomalous diffusion coefficient, *t* is time, *α* is the anomaly parameter and *c*_1_ is the error in detection. The *(b)* diffusion with drift model is:

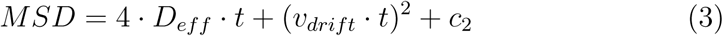

where *D*_*eff*_ is the effective diffusion coefficient, *t* is time, *v*_*drift*_ is the drift velocity and *c*_2_ is the error in detection. The data is fit to 3/4th of the MSD trajectory to avoid artifacts due to small sample sizes for very large time points (46, 61).

### Calibrating motor density

Affinity purified 6x-His tagged EGFP was used as a standard to calibrate GFP-tagged dynein concentration. 50 *μ*l EGFP was flowed into the flow chamber in a concentration series ranging from 1 ng/*μ*l to 100 ng/*μ*l and imaged in a fluorescence microscope. A 3D stack of images was acquired every 4 *μ*m steps with an exposure time of 150 ms. Average intensity projections from five different fields of view were used to estimate the intensity per unit area. The point spread function (PSF) of the lens was estimated from a z-scan of 0.2 *μ*m FITC labelled fluorescent beads (Invitrogen, ThermoFisher Scientific). The intensity profile through the bead centre was fit to a gaussian function:

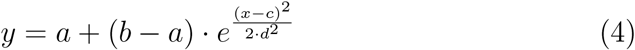

where a and b are scaling factors, c is the mean (*μ*) and d is the standard deviation (*σ*) using the *CurveFitter* function in ImageJ (Figure 1(A)-(C)). Using the full width at half maximum FWHM= 1.84 *μm* (≈ 0.23548*σ*) the volume of every image and the number of molecules in the field of view were estimated. The GFP-tagged dynein motor concentration (*c*_*m*_) was estimated from the fit to the intensity of an EGFP calibration curve (Figure S1 (D)). A conversion factor of 2.01:1 was used to compensate for the relative brightness of EGFP compared to GFP (62) and the linear motor density 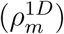 is calculated as 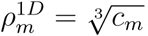, assuming a uniform distribution of motors. The number of motors acting on individual filaments in experiment was then simply estimated from the MT length (*L*) and motor-density per unit length 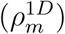 as follows: 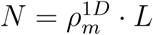.

### Model

The model of microtubule and motor mechanics is based on previous reports (34, 63–67). Specific aspects that were modified from previous work are described in the following sections. Parameters were taken from previous reports of experimental measurements, where available, and in their absence reasonable estimates were made (Table 1).

**Table 1.**
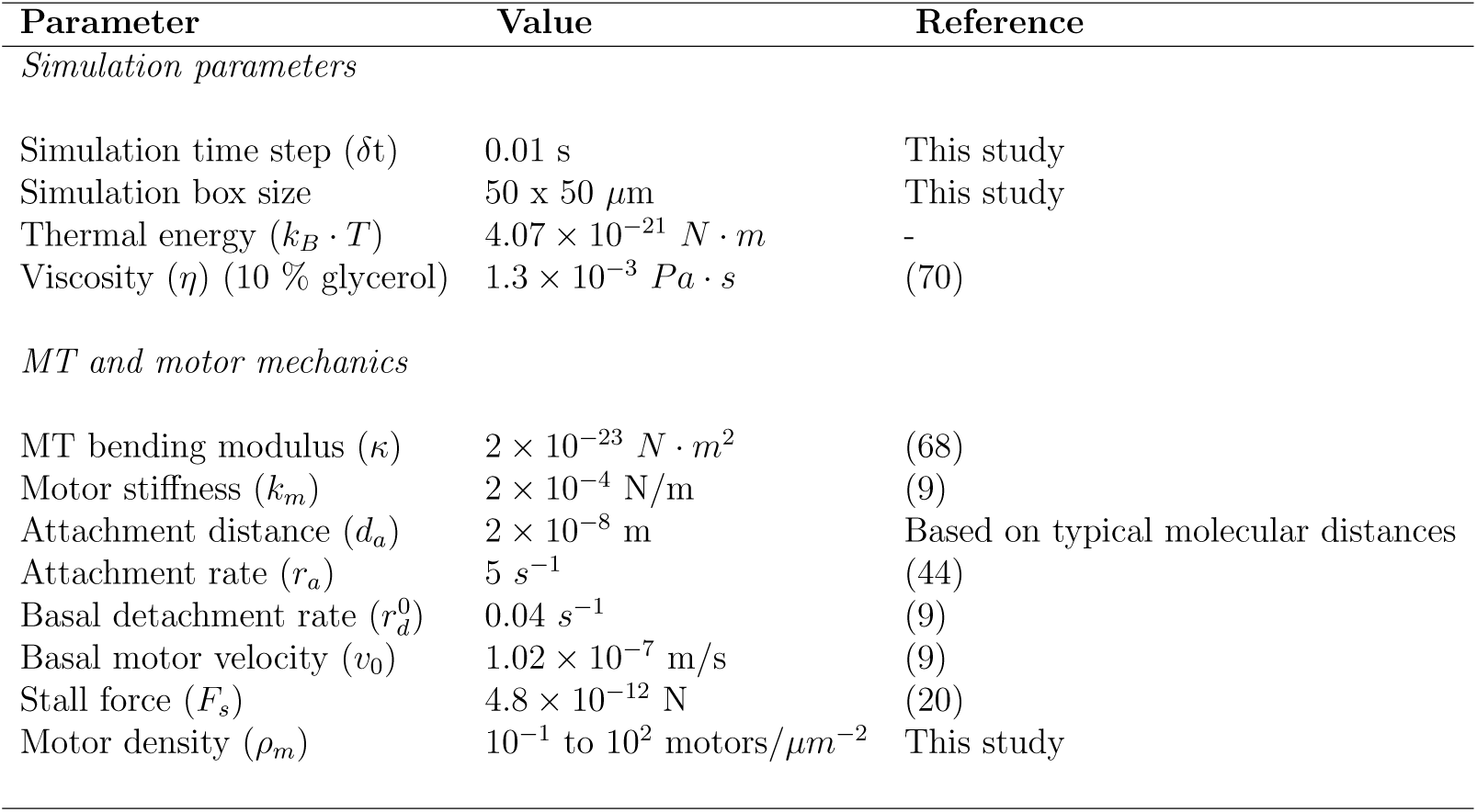
Simulation parameters of the gliding assay. Parameters of the mechanics of MTs and dynein are taken from literature while the motor density was varied.

| Parameter | Value | Reference |
| --- | --- | --- |
| *Simulation parameters* |  |  |
| Simulation time step ($\delta t$) | 0.01 s | This study |
| Simulation box size | $50 \times 50 \mu\text{m}$ | This study |
| Thermal energy ($k_B \cdot T$) | $4.07 \times 10^{-21} \text{ N} \cdot \text{m}$ | - |
| Viscosity ($\eta$) (10 % glycerol) | $1.3 \times 10^{-3} \text{ Pa} \cdot \text{s}$ | (70) |
| *MT and motor mechanics* |  |  |
| MT bending modulus ($\kappa$) | $2 \times 10^{-23} \text{ N} \cdot \text{m}^2$ | (68) |
| Motor stiffness ($k_m$) | $2 \times 10^{-4} \text{ N/m}$ | (9) |
| Attachment distance ($d_a$) | $2 \times 10^{-8} \text{ m}$ | Based on typical molecular distances |
| Attachment rate ($r_a$) | $5 \text{ s}^{-1}$ | (44) |
| Basal detachment rate ($r_d^0$) | $0.04 \text{ s}^{-1}$ | (9) |
| Basal motor velocity ($v_0$) | $1.02 \times 10^{-7} \text{ m/s}$ | (9) |
| Stall force ($F_s$) | $4.8 \times 10^{-12} \text{ N}$ | (20) |
| Motor density ($\rho_m$) | $10^{-1} \text{ to } 10^2 \text{ motors}/\mu\text{m}^{-2}$ | This study |

### Microtubules

MTs were modeled as polymers of a constant length (L) and with a bending modulus (*κ*) reported previously (68), to recapitulate experimental conditions. For visualization, the values of L were drawn from an exponential probability distribution (Equation 1) based on parameters that were obtained from fitting to experimental data (Figure S3). As filaments move through the fluid, they experience a drag force (*F*_*d*_). Assuming a cylindrical geometry and simplifying the drag term as an average between the longitudinal and transverse components (a two-fold difference), as previously described by (64), the mobility is given by: *μ* = log (*L/δ*)*/*3 · *π · η · L*, where L is the MT length, *δ* is the diameter of the rod and *η* is the viscosity of the fluid. This results in *F*_*d*_ = *v/μ*, where *v* is the velocity of the transported MT.

### Motor model

The motors are immobilized by their tails on the surface, and the heads can bind, unbind and step along the filament. The motor density (*ρ*_*m*_) was varied between 0.1 and 100 motors/*μ*m^2^ including 2 and 16 motors/*μ*m^2^ based on experimental estimates. The details of the motor attachment, detachment and stepping behavior that determine the transport characteristics, are described in the following sections.

*Attachment and detachment:* Motor heads bind stochastically to MTs based on an attachment rate (*r*_*a*_) if they are within a distance of attachment (*d*_*a*_) to the filament (Figure 1). Motor detachment was modeled to be probabilistic, load dependent and anisotropic, i.e. different for forward and backward load forces based on the single molecule unbinding measurements of the yeast dynein (43). The precise function defining the load dependence of *r*_*d*_ was inferred by digitizing the reported values (43) using *Plot Digitizer* (http://plotdigitizer.sourceforge.net/) and fitting them to a custom piece wise function:

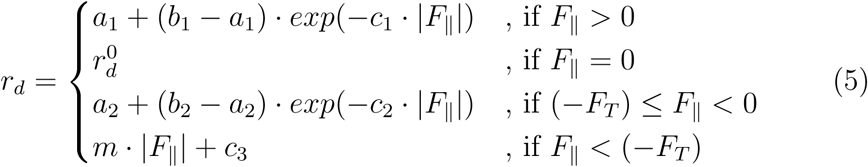

*where, F*_*||*_ corresponds to the component of motor extension along the microtubule and *F*_*T*_ is a threshold force magnitude with the positive (+) sign indicating backward and negative (-) sign indicating forward loads (Figure S6). *F*_*T*_ = 3.51728 pN is the threshold force inferred from the fit. The other fit parameters for the backward load profile (Goodness of fit R^2^ = 0.67) are *a*_1_ = 2.3159, *b*_1_ = 0.0400, *c*_1_ = 0.8414. The two-part fit to the forward load profile has parameters (i) (Goodness of fit R^2^ = 0.90) *a*_2_ = 7.2768, *b*_2_ = 0.0400, *c*_2_ = 0.6426 for the first part and (ii) (goodness of fit R^2^ = 1) m = 3.5814, *c*_3_ = −6.4016 for the second part. All fitting was performed using the MATLAB Optimization Toolbox (Mathworks Inc., USA).

*Motor stepping:* Based on single molecule reports discrete step-sizes and the force dependence of dynein motility (9, 20), we have implemented a discrete stepping model of motor movement. The motors step probabilistically at a constant stepping rate of 8 *s*^*-*1^ (69). The instantaneous step size is modeled to be load-dependent based on the following relation:

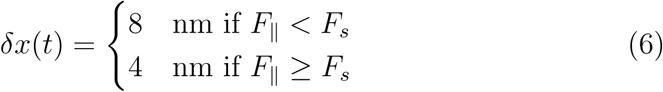

where *F*_*||*_ is the load force parallel to the MT to which the motor is bound and *F*_*s*_ is the stall force. The force-dependent, stochastic forward or backward stepping is modeled by a probability of forward stepping (*p*_*f*_) based on multiple force thresholds (*f*_1_ to *f*_4_) obtained from the full length yeast dynein (20) and scaled for the stall force of the truncated construct, as follows:

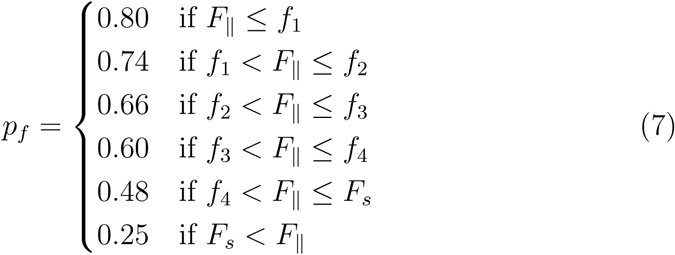

where *f*_1_ = 0 pN, *f*_2_ = 1 pN, *f*_3_ = 3 pN and *f*_4_ = 4 pN. The probability of backward (*p*_*b*_) stepping is *p*_*b*_ = 1 ‒ *p*_*f*_.

### Simulations

2D simulations were performed using Cytosim, a C++ based Langevin dynamics simulation engine (64). Space was modeled as a square simulation box with periodic boundary conditions. The integration time was chosen to be smaller than the fastest time-scale (Table 1). The system consists of MT filaments of a fixed length (*L*) and a fixed number of motors determined by the density (*ρ*_*m*_). Both filaments and motors were randomly distributed in simulation space at the time of initialization.

## Author Contributions

KJ performed experiments and analyzed data; NK designed model and simulations; CAA designed the research and wrote the manuscript.

## Acknowledgments

We would like to thank Ron Vale and Andrew Carter for the gift of the *S. cerevisiae* VY208 strain and Thomas Pucadyil for the purified EGFP and TEV protease. A grant from the Dept. of Biotechnology, Govt. of India (BT/PR6715/GBD/27/463/2012) and IISER Pune core-funding to CAA supported the work. KJ and NK were supported by fellowships from DST-INSPIRE (IF-130394) and CSIR (09/936(0128)/2015-EMR-1) respectively.

---

**Figure S1:**
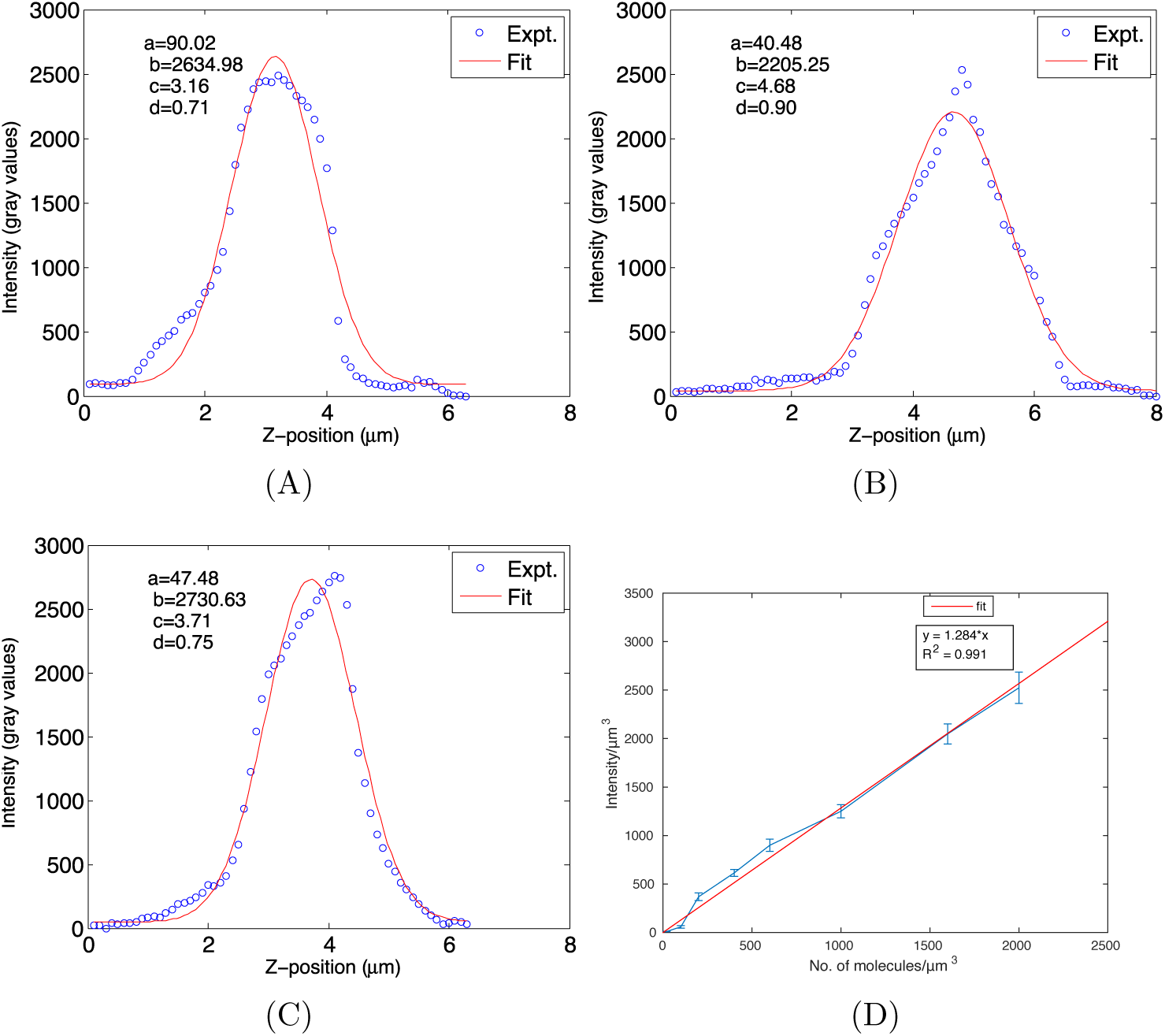
PSF estimation. (A)-(C) The grey value intensity profiles (blue circles) of three representative fluorescent beads of diameter 200 nm are plotted through the z-slices (1 slice = 100 nm) through the XY centre of the bead. The profiles are fit to a gaussian function (red line). The fit parameters are used to estimate the FWHM as described in the methods section. (D) Images of EGFP in the motility flow chamber are used to plot mean (*±* s.d.) intensity in grey values per *μ*m^3^ (y-axis) as a function of number of molecules per *μ*m^3^ (x-axis) (blue). The linear fit (*R*^2^ = 0.99) to the intensity profile, *y* = 1.284 · *x* (red line) is used to estimate the motor density from the unknown sample.

**Figure S2:**
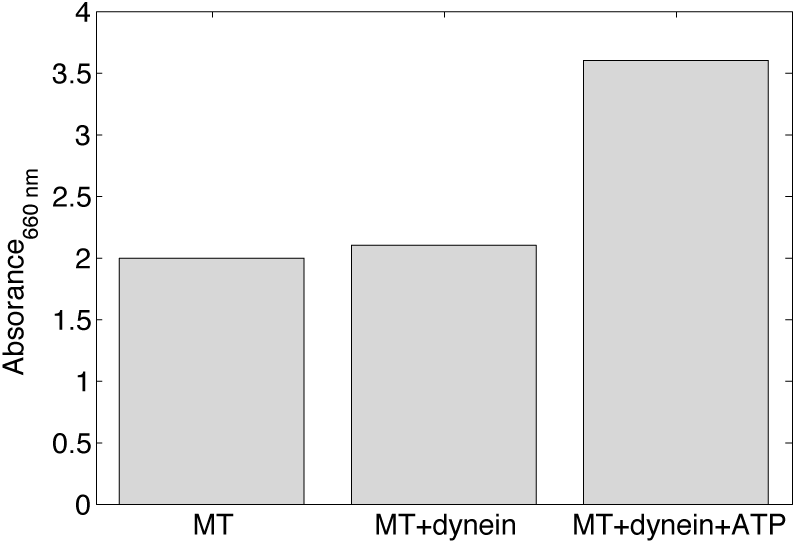
Testing dynein activity. The activity of the purified dynein was tested by measuring the absorbance at 660 nm (y-axis) for samples containing MTs alone, MTs with dynein and MT with dynein and 1 mM ATP (x-axis) in a malachite green assay for inorganic phosphate release.

**Figure S3:**
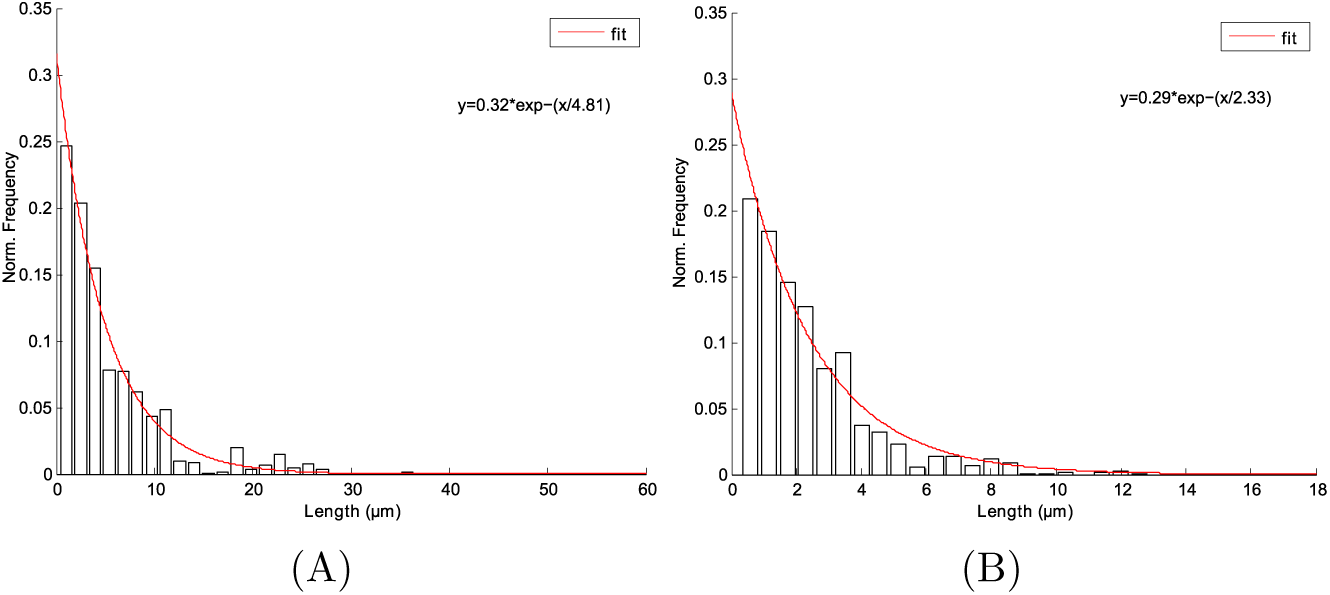
MT length distributions. Taxol stabilized MT filament lengths were analyzed and the sum normalized frequency distributions of filaments incubated for 24 hours (A) before and (B) after shearing (as described in the Methods) are plotted. The sum normalized distributions are fit to an exponential decay function *y* = *A · e*^*-L/λ*^ (red curve), where *A* is the scaling factor and *λ* is the mean length of the population.

**Figure S4:**
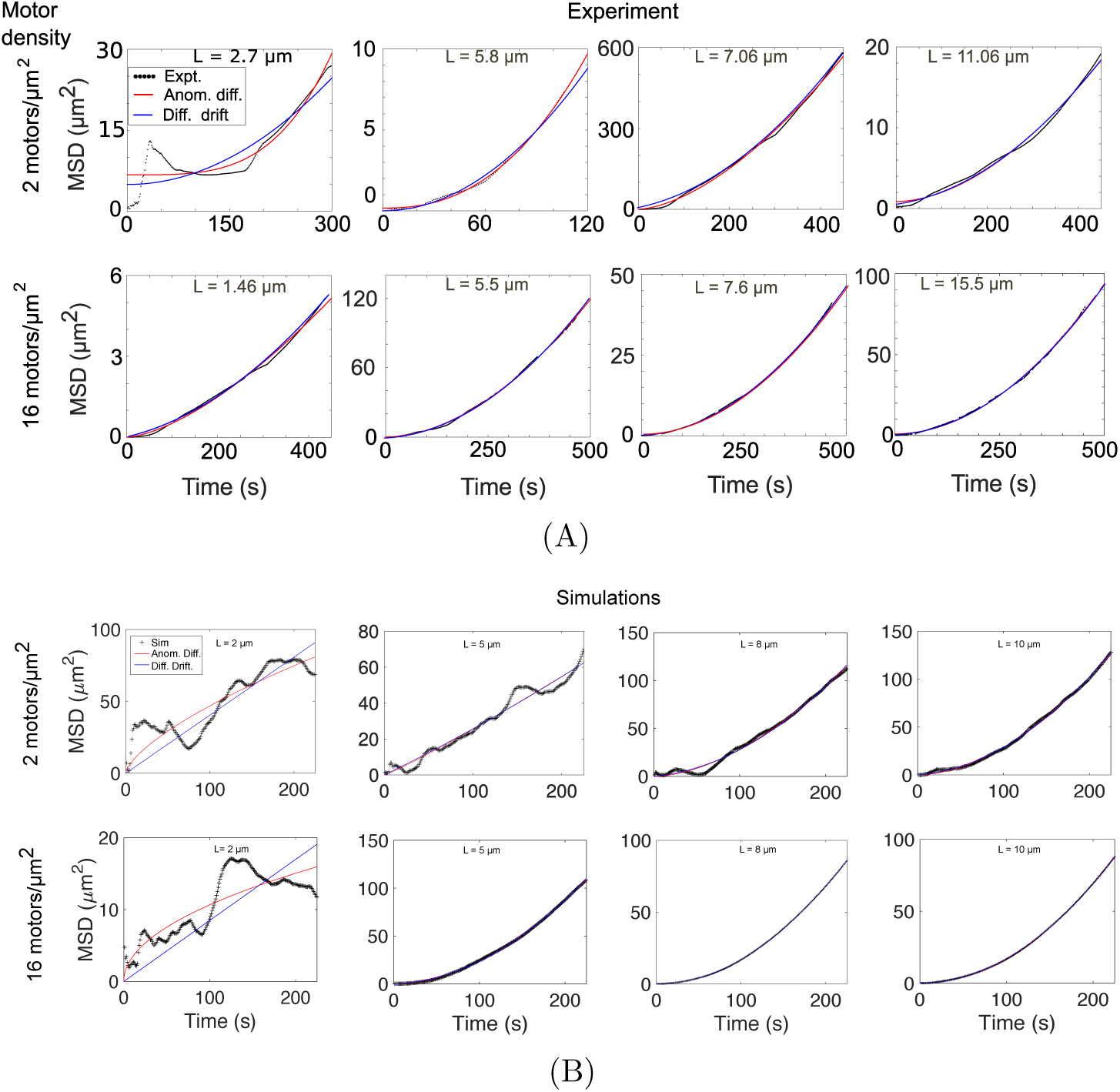
MT length dependence of Mean square displacement (MSD) from experiments and simulation. (A) Experimentally measured MT transport was quantified in terms of MSD as a function of time (black) for increasing MT filament lengths (L). Representative plots of MTs of length ranging from 1.46 to 15.5 *μ*m when motor density was *ρ*_*m*_ = 2 (upper panel) and *ρ*_*m*_ = 16 motors/*μ*m^2^ (lower panel) are plotted. (B) Simulation based MSD profiles in presence of the same range of *ρ*_*m*_ as above, are plotted representatively for MTs of lengths 2, 5, 8 and 10 *μ*m. Both experimental and simulated MSD profiles were fit to anomalous diffusion (red, Equation 2) and diffusion with drift models (blue, Equation 3).

**Figure S5:**
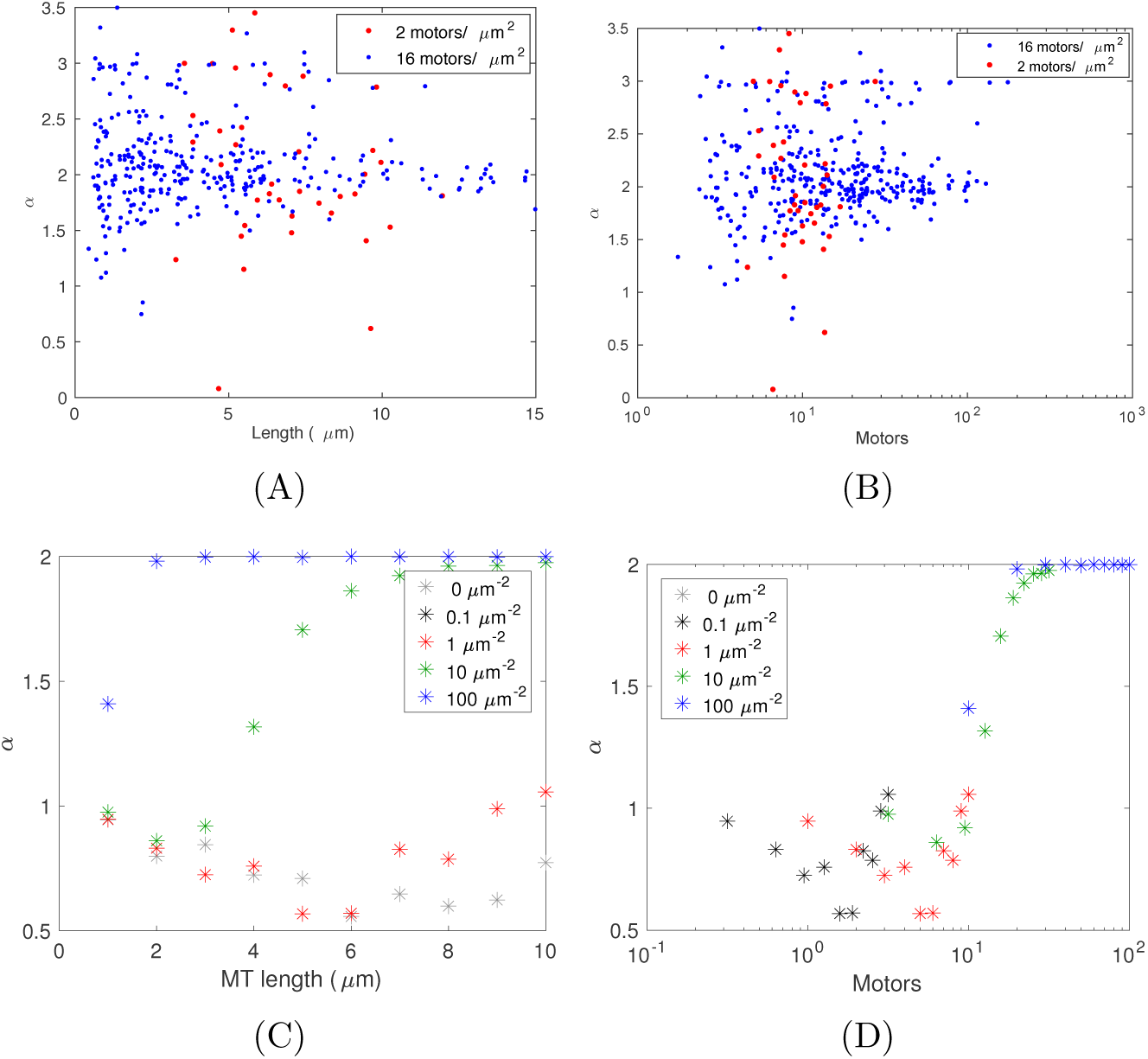
Anomaly parameters from experiment and simulation. (A)-(B) The experimental estimates of the anomaly parameter of diffusion (*α*) is plotted as a function of (A) MT length and motors and compared to (C)-(D) estimates from simulation as a function of (C) MT length and (D) number of motors. The anomaly parameter *α* is obtained by fitting Equation 2 to the MSD plots (Figure S4). The colors indicate the different motor densities.

**Figure S6:**
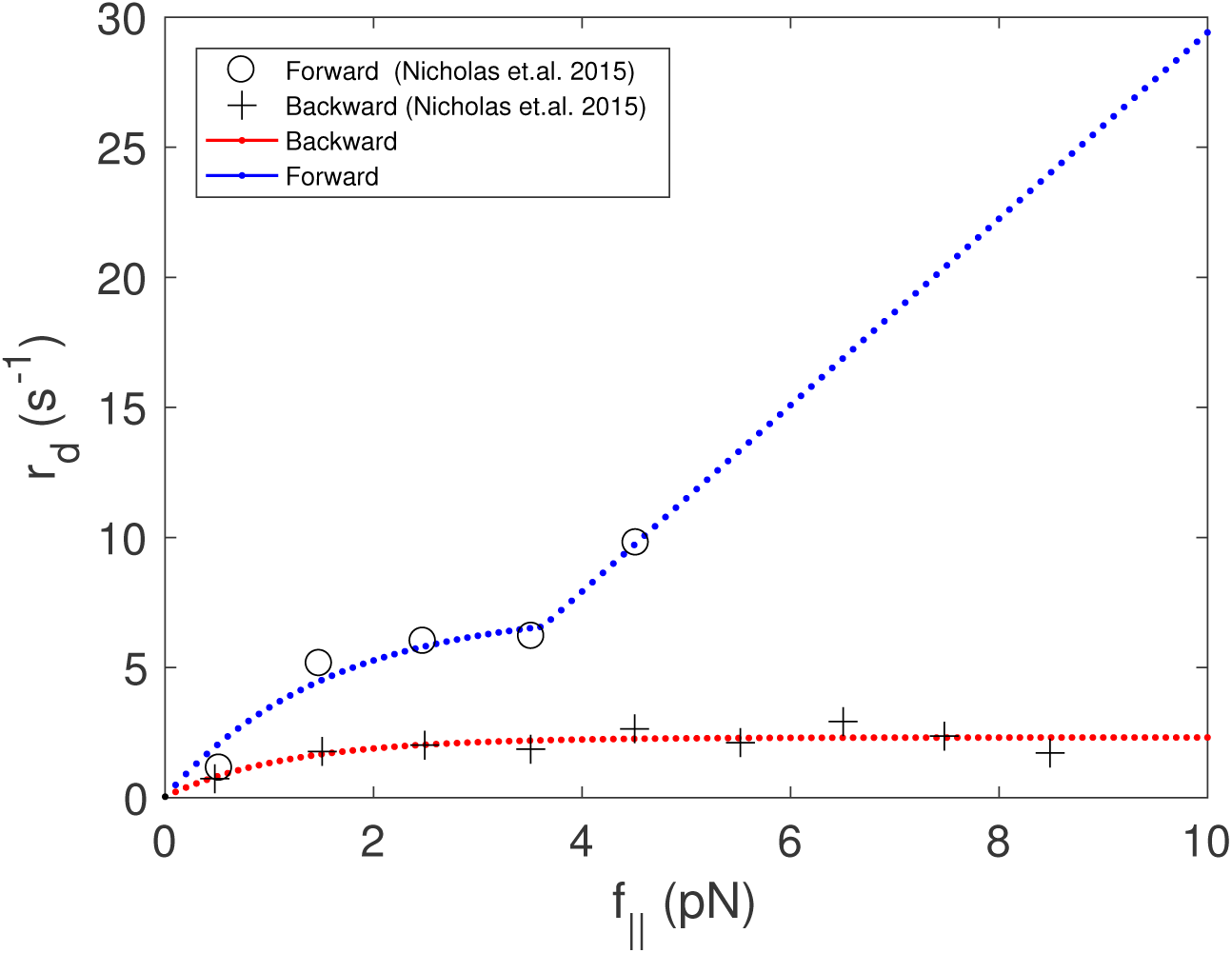
Anisotropic detachment rate model. The experimental data from single molecule experiments (43) were digitized to plot the measured *r*_*d*_ under forward (o) and backward (+) loads. The data was then fit by a custom function (Equation 5) for the rate of detachment, depending on forward (blue) and backward (red) load forces.

**Figure S7:**
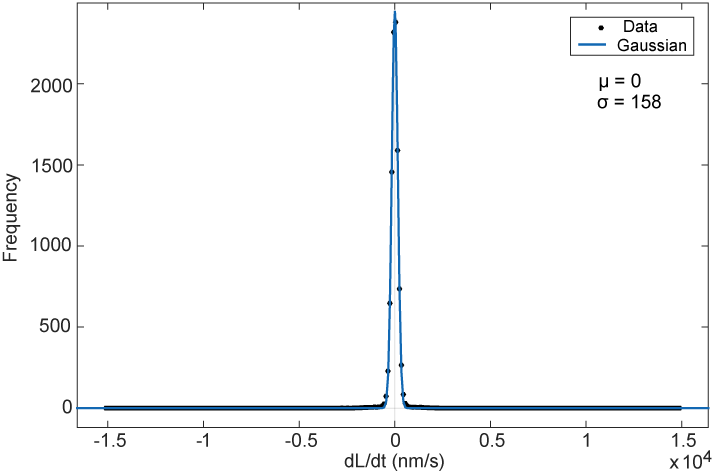
Estimating MT length variations. MTs were tracked using the nanometer precision filament tracking program, FIESTA (45), and the frequency distribution of the pairwise change in length as a function of time (*dL/dt*) in nm/s (black circles) was fit to a gaussian (blue curve) with a mean of 0 nm/s and standard deviation of 158 nm/s.

**Figure S8:**
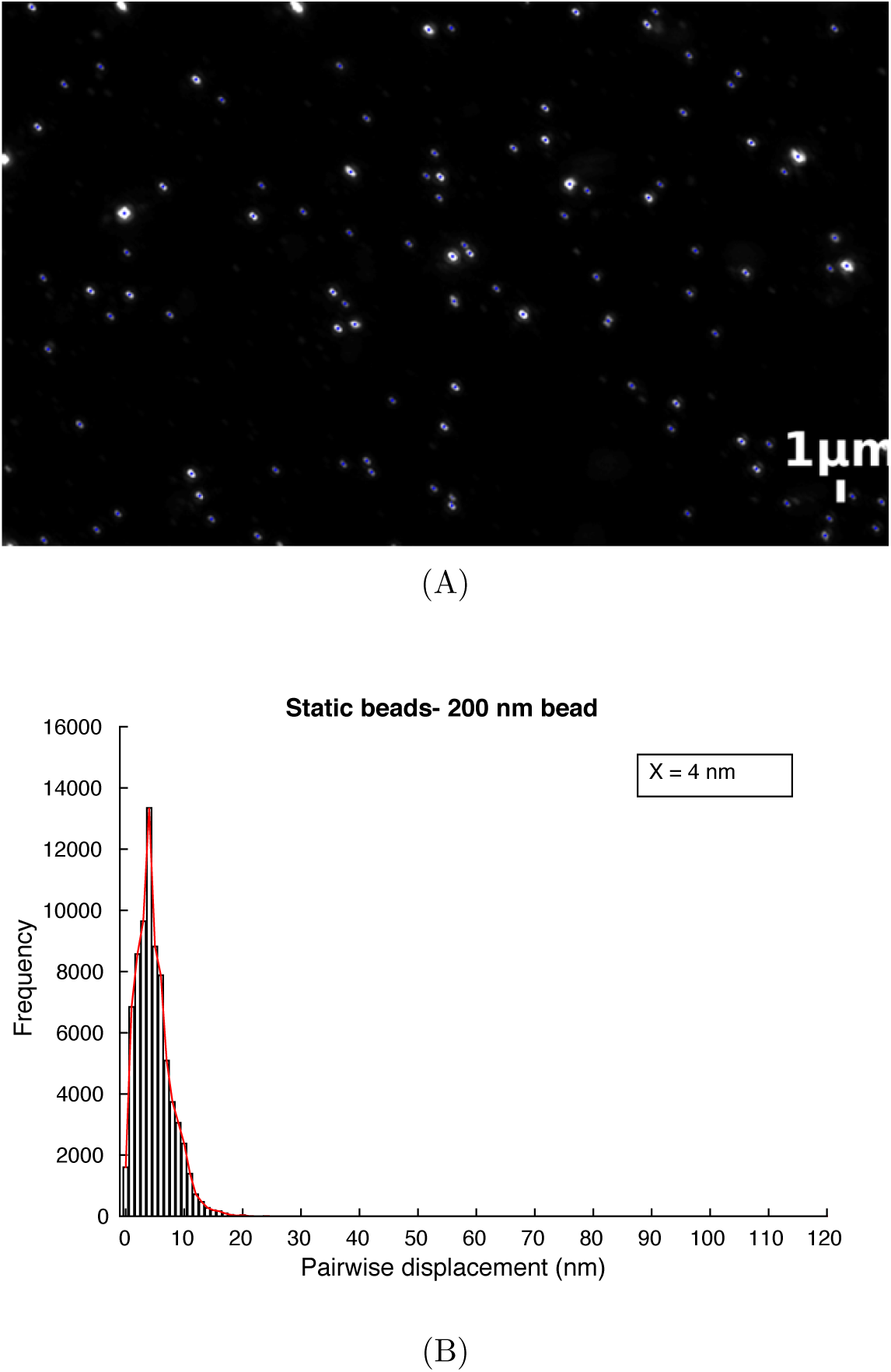
Mobility of fixed beads. (A) A time-series of fluorescently labelled 0.2 *μ*m beads immobilized on a glass slide was tracked (blue lines) and (B) the pairwise displacement histogram (black bars) was fit (red line) using a peak detection algorithm in MATLAB and showed a peak at 4 nm.

## Supplemental movies

**Video 1:**
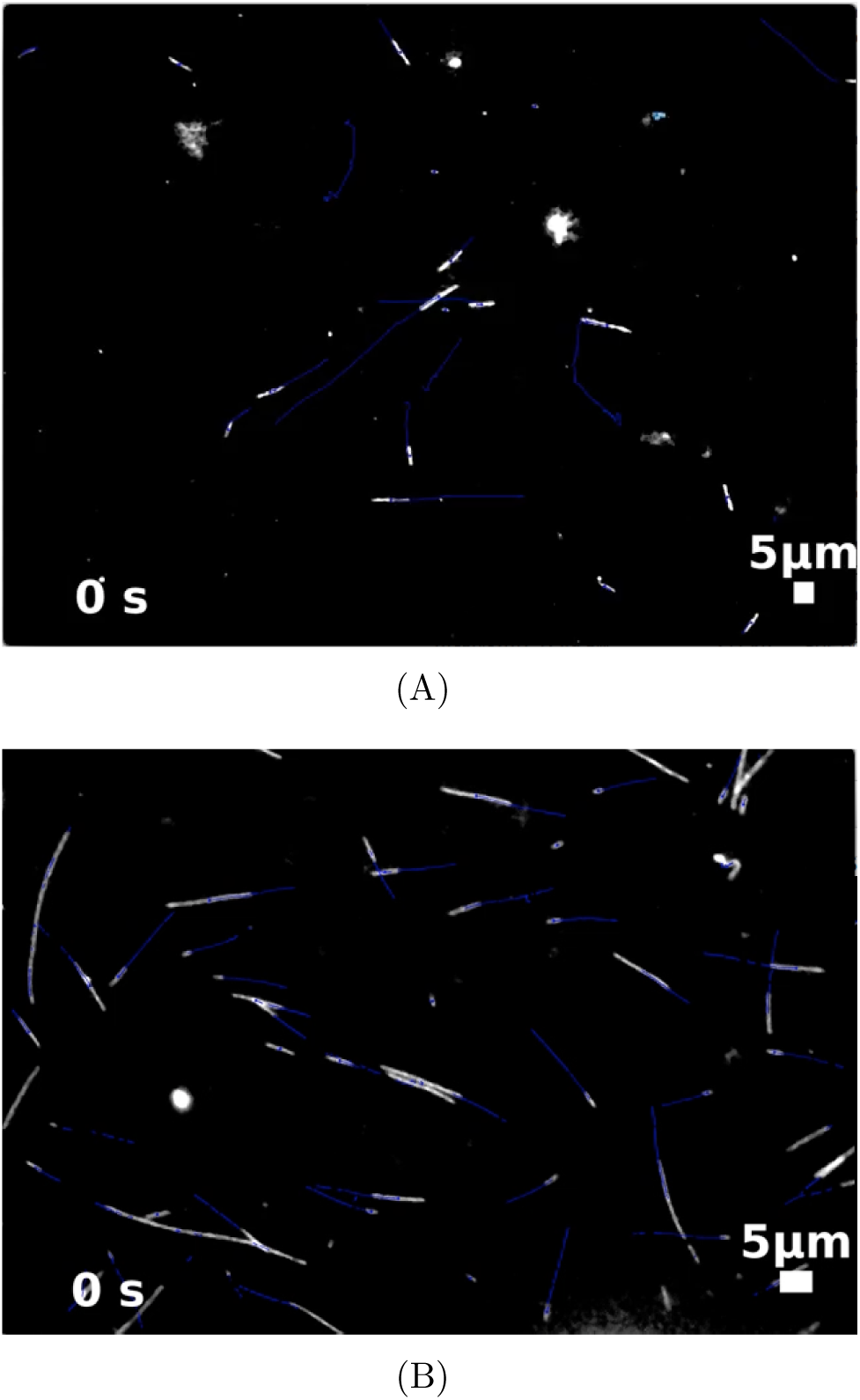
Time-lapse movies of MT gliding in experiment. A time series of MTs gliding in the presence of (A) 2 and (B) 16 motors/*μ*m^2^ dynein were acquired every 1 second. The rhodamine labelled taxol-stabilized MTs (grey) were tracked using FIESTA and tracks are overlaid on the time series (blue lines). The time stamp is in units of seconds and the scale bar is 5 *μ*m.

**Video 2:**
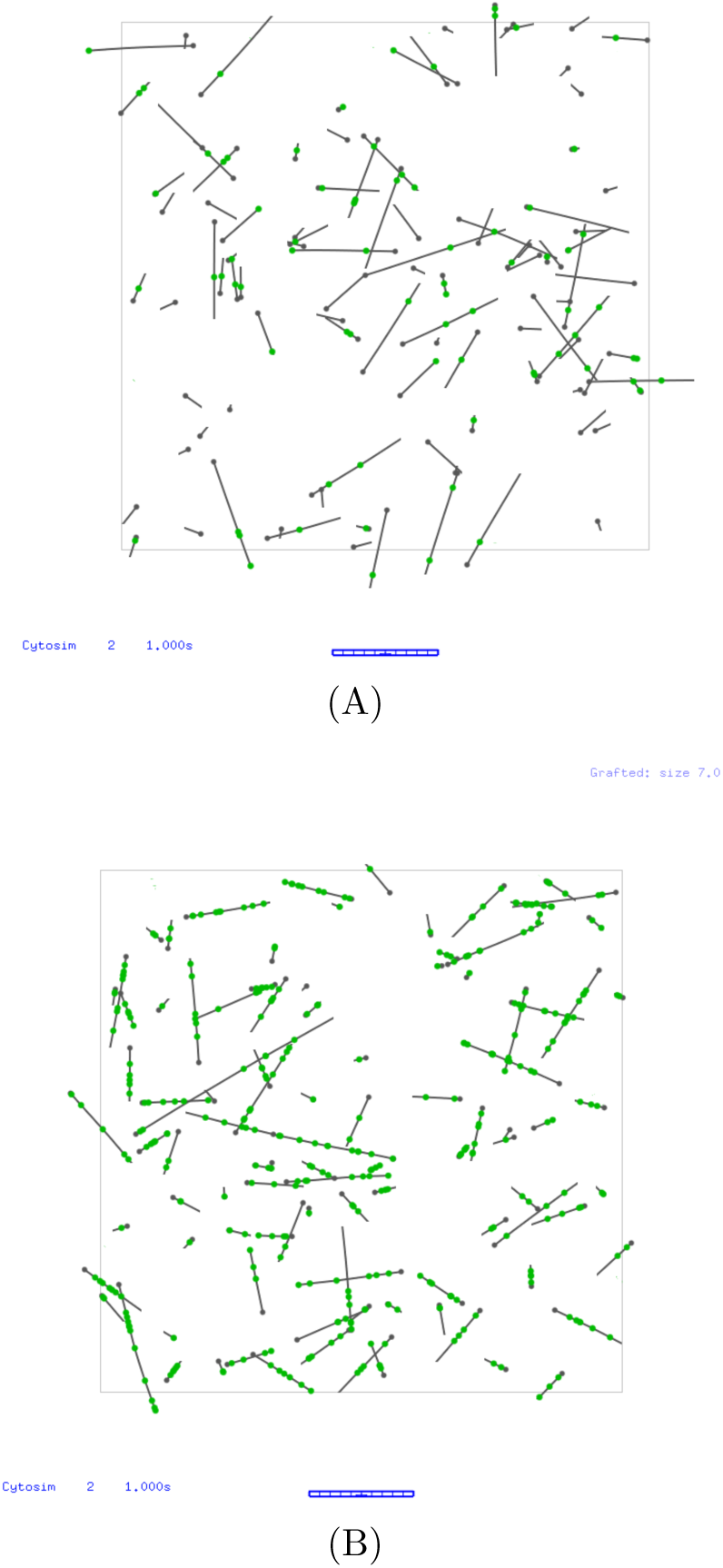
Simulations of 2D MT transport by dynein. Simulations of semi-flexible polymers of microtubules as they glide and swivel when attached to motors, and freely diffuse if they are unbound, were performed using Cytosim, with motor densities (A) 2 and (B) 16 motors/*μ*m^2^. Individual MT lengths are constant during the simulation to mimic taxol-stabilized filaments and exponentially distributed to reproduce experimental observations. Scale bar 10 *μ*m.

